# DNCB shows hormetic effects in THP-1 cells: low-dose enhancement of metabolic activity

**DOI:** 10.64898/2026.08.19.745690

**Authors:** Darja Henseler, Osmond Aruna

## Abstract

2,4-Dinitrochlorobenzene (DNCB) is a well-characterized skin sensitizer that has been widely used in immunological and toxicological research and, historically, in clinical immunotherapy. Although it is a well-investigated chemical, this is the first study focusing on the dose–response behavior at low-level concentrations. The aim was to reveal potential hormetic effects due to its known Nrf2 inducting activity. Therefore, THP-1 cells were treated with low doses of DNCB and two endpoints were evaluated for hormetic responses: metabolic activity using a resazurin-based assay and immune activation by measuring CD86 and CD54 expression using flow cytometry. The results showed a significant hormetic effect on the metabolic endpoint at the lower cell density for both analyzed time points, and a hormetic tendency at the higher cell density. Metabolic activity increased to approximately 125% of the control at 0.05 µM DNCB. For the immunological endpoint a slight decrease in CD86 and CD54 surface marker expression was observed, up to −16% and up to −12% compared to control at 0.5 µM DNCB. These findings highlight the importance of including low-dose concentrations when characterizing chemical dose-response relationships and evaluating toxicological risk.

## Introduction

Hormesis describes a dose-response phenomenon in which low doses of stressors or harmful substances can trigger a positive, adaptive response in biological systems. This biphasic behavior is often reflected in a U-shaped or J-shaped dose-response curve, with low doses having a stimulating effect and high doses having a toxic effect. The earliest publications on this phenomenon can be traced back to the work of Hugo Schulz. Around 1880, Schulz demonstrated that when yeast was exposed to various toxic substances, low doses of these substances stimulated fermentation activity, whereas higher doses reduced or completely inhibited fermentation (1). Biphasic dose-response relationships have been observed in a wide range of organisms including humans, animals, plants, microorganisms and even to single cells. To characterize hormetic dose–response relationships, Calabrese et al., 2011 developed a database comprising approximately 9,000 biphasic dose–response cases and analyzed the hormetic ranges and the maximum stimulation levels. The analysis showed that stimulation in the low-dose range is generally moderate, with most cases exhibiting maximum stimulation between 130% and 160% of the control, corresponding to a 30% to 60% increase over baseline. The hormetic effect range is usually 10 to 20 times below the lowest dose for a toxicological or pharmacological effect (2). The endpoints for which a hormetic dose-response relationship can be demonstrated are considered highly integrated and include, for example, growth, metabolic activity, immune response, survival, neurological, cancer, and other endpoints (2, 3). The most frequently investigated endpoints in animals were metabolic activity (24%) and growth (19%) (2). The mechanisms underlying hormesis generally involve the stimulation of cellular processes that are critical for maintaining homeostasis (4). A key mechanism through which hormetic effects are mediated is the activation of the transcription factor Nuclear factor erythroid 2-related factor 2 (Nrf2) (5). Nrf2 regulates diverse cellular processes such as differentiation, proliferation, and inflammation by activating antioxidative gene expression while inhibiting reactive oxygen species production, proinflammatory cytokines, and NF-κB signaling. Aberrant Nrf2 function or expression is linked to pathologies including inflammatory diseases, autoimmunity, cancer, and neurodegeneration (6, 7). For many secondary plant compounds, acting as Nrf2 inducers, a hormetic effect has already been demonstrated (8). In this study, we aim to investigate whether the synthetic compound 2,4-dinitrochlorobenzene (DNCB), which also activates the Nrf2 pathway (9-11), also exhibits hormetic effects at the cellular level. DNCB serves in industrial production as a starting material for explosives, dyes, rubber auxiliaries, and photochemicals and is also used in research to induce controlled sensitizations in immunological and toxicological studies, due to its sensitizing potential. Additionally, in some cases, DNCB is used together with a chemotherapeutic agent for the treatment of cutaneous metastases (12, 13). In this context, the local DNCB-mediated immune reaction is utilized, which enhances antitumor effects.

In our study, we first address the most frequently analyzed endpoint in hormetic studies, which is a metabolic activity, assessed by using a resazurin-based assay. Second, given that DNCB is a well-known contact allergen, we examined potential hormetic effects on immunological endpoints in the human monocytic leukemia cell line THP-1, a commonly used model for chemical sensitization, by quantifying the upregulation of the costimulatory surface protein CD86 and the adhesion protein CD54.

## Results

### Determination of optimal cell density based on assay linearity

A calibration curve was established using varying numbers of THP-1 cells per well, ranging from 10,000 to 220,000 cells in a total volume of 180 µL RPMI medium (corresponding to 0.056 × 10^6^ to 1.22 × 10^6^ cells/mL). At 2 h and 24 h post-seeding, cells were incubated with PrestoBlue™ HS cell viability reagent, and fluorescence intensity (FI) was measured. PrestoBlue™ assays are typically performed using low cell numbers to ensure measurements remain within the linear detection range. The data showed a linearity between cell number of 10,000 up to 220,000 cells. At this density, a 23.6-fold (2 h) and 21.6-fold (24 h) increase in FI was observed, which closely corresponds to the theoretically expected 22-fold increase relative to 10,000 cells (Table 1). As lower cell numbers are generally preferred for viability assays, the lowest tested cell density exhibiting a robust signal-to-blank ratio (≥5-fold above background) was selected. Accordingly, 80,000 cells (0.44 × 10^6^ cells/mL) were used for subsequent experiments. Given that previous studies (13) have demonstrated that hormetic responses may depend on experimental conditions, both multiple time points and cell densities were evaluated. In addition, we included a cell density of 1.0 × 10^6^ cells/mL (180,000 cells per well), modified from the original h-CLAT protocol (160,000 cells per well). For subsequent experiments, the 2 h time point was replaced by 4 h to align with commonly used conditions for assessing early metabolic responses.

**Table 1.** Determination of optimal cell density based on fluorescence linearity. Fluorescence intensities (FI) of varying THP-1 cell numbers measured at 2 h and 24 h post-seeding are shown. In addition, expected FI ratios and experimentally observed FI ratios between cell densities are presented, demonstrating a near-linear relationship across the tested range.

| Cells per well | FI (2 h) | FI (24 h) | Expected ratio | ratio FI (2 h) | ratio FI (24 h) |
| --- | --- | --- | --- | --- | --- |
| 10,000 | 166 | 310 | 1 | 1.0 | 1.0 |
| 40,000 | 666 | 1355 | 4 | 4.0 | 4.4 |
| 80,000 | 1334 | 2936 | 8 | 8.0 | 9.5 |
| 100,000 | 1583 | 3562 | 10 | 9.5 | 11.5 |
| 140,000 | 2293 | 4738 | 14 | 13.8 | 15.3 |
| 180,000 | 2810 | 5566 | 18 | 16.9 | 18.0 |
| 220,000 | 3920 | 6695 | 22 | 23.6 | 21.6 |

### DNCB induces hormetic effects on metabolic activity in THP-1 cells

To assess whether DNCB induces hormetic effects on metabolic activity, THP-1 cells (80,000 cells/well and 180,000 cells/well) were treated with seven concentrations of DNCB, ranging from 0.05 µM to 40 µM, for 4 h and 24 h. A significant hormetic response was detected at 80,000 cells. At 180,000 cells, the dose– response curve displayed a hormetic shape, but the effect was not statistically significant and is therefore described as a hormetic tendency (see Figure 1). Specifically, a maximal increase in metabolic activity to 125% was observed at 0.05 µM DNCB for 80,000 cells/well (4 h and 24 h), which was statistically significant at both time points (p<0.05). For 180,000 cells/well the maximum increase was lower, with metabolic activity reaching 115% (4 h) and 112% (24 h) at 0.05 µM DNCB.

**Figure 1.**
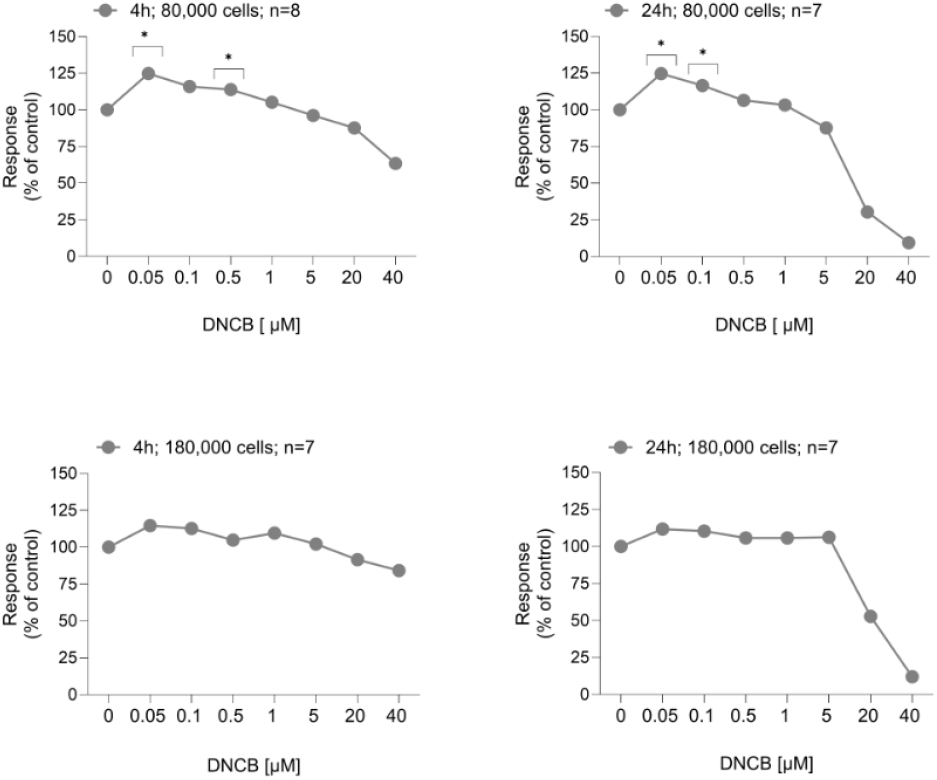
2,4-Dinitrochlorobenzene (DNCB) induces a hormetic response in THP-1 cells. THP-1 cells (80,000 and 180,000 cells/well) were treated with DNCB ranging from 0.05 µM to 40 µM for 4 h or 24 h. Metabolic activity is presented as percentage relative to the control (DMSO 0.2%), demonstrating increased responses between 0.05 µM and 0.5 µM DNCB, with statistically significant increases (p<0.05) at 0.05 µM and 0.5 µM for 80,000 cells (4 h), at 0.05 µM and 0.1 µM for 80,000 cells (24 h). For the evaluation of significance, only the dose range between 0.05 µM and 1 µM was considered, as we wanted to focus on the low dose range.

For 80,000 cells at 0.1 µM, metabolic activity increased to 116% (4 h) and 117% (24 h), with statistical significance observed at 24 h (p<0.05). At 0.5 µM, metabolic activity increased to 114% (4 h) and 106% (24 h), reaching statistical significance at 4 h (p<0.05). At 1 µM DNCB, only a slight, non-significant increase in metabolic activity was observed, not reaching statistical significance. Thus, the hormetic response was primarily observed between 0.05 µM and 0.5 µM DNCB, corresponding to a stimulation range spanning one order of magnitude. In addition, a decline in metabolic activity was observed at concentrations ≥ 20 µM DNCB. This cytotoxic effect was more pronounced at lower cell densities compared to higher cell densities, and at longer exposure times compared to shorter incubation periods.

### Effects of low-dose DNCB on surface marker expression of CD86 and CD54

DNCB is known as a strong sensitizer used as positive control in the human cell line activation test (OECD Test Guideline 442E) by analysing upregulating of THP-1 cell surface markers CD86 and CD54. As the strong sensitization effect of 20 µM DNCB is known and even for lower doses as 5 µM an upregulation was reported (14) we were interested in investigating the low-dose effect of DNCB on the immunological endpoint by analysing surface marker expression of CD86 and CD54. Therefore, THP-1 cells were exposed to DNCB (0.5 µM to 40 µM) for 24 h. Following surface marker expression of CD86 and CD54 was analysed by flow cytometry.

First, we analyzed 180,000 cells per well, a cell number selected based on the cell numbers used in the human cell line activation test. Consistent with the literature, a pronounced upregulation of CD86 (293%) and CD54 (477%) was observed at 20 µM DNCB compared to control, while mean cell viability was 73%. In experiments with 80,000 cells per well, CD86 and especially CD54 showed a shifted and lower upregulation. Upregulation was already detectable at 5 µM DNCB for CD86 (128%) and for CD54 (170%) and remained at 20 µM DNCB (CD86: 128%; CD54: 170%), with cell viability of 95% at 5 µM DNCB and 72% at 20 µM DNCB.

Interestingly, in the low-dose range, a decreased expression of CD86 and CD54 was observed. Experiments with 180,000 cells per well showed a decrease in CD86 at 0.1 µM DNCB (−8%), at 0.5 µM DNCB (−13%), and at 1 µM DNCB (−16%), while experiments with 80,000 cells per well resulted in very similar values at 0.1 µM (−10%), at 0.5 µM (−16%), and at 1 µM (−16%). This concentration range overlaps with the hormetic range identified for metabolic activity. For CD54 expression, an additional shift was observed between the two cell densities. CD54 showed decreased expression between 0.5 µM DNCB (−9%) and 5 µM DNCB (−12%) for 180,000 cells per well, and between 0.1 µM DNCB (−7%) and 1 µM DNCB (−12%) for 80,000 cells per well. Cell viability in the low-dose range of DNCB was comparable to the control, as assessed by propidium iodide staining (100 ± 1%).

In summary, low concentrations of DNCB exerted a slight inhibitory tendency on CD86 and CD54 expression (see Figure 2 A+B).

**Figure 2.**
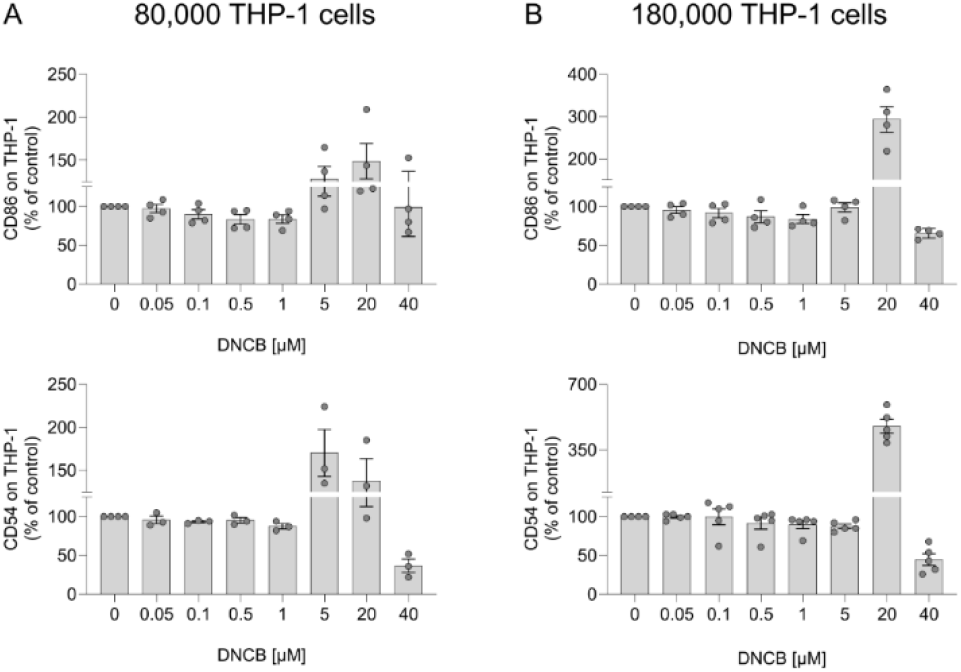
Effect of low-dose 2,4-dinitrochlorobenzene (DNCB) on surface expression of CD86 and CD54. A) 80,000 THP-1 cells/well and B) 180,000 THP-1 cells/well were treated with DNCB (0.05 µM to 40 µM) for 24 h and stained with fluorochrome-conjugated antibodies against CD86 and CD54. Relative fluorescence intensity (RFI), expressed as percentage of control, is shown. A slight decrease in expression was observed at low DNCB concentrations for CD86 (up to −16%) and CD54 (up to −12%).

## Discussion

Hormetic dose-response relationships are ubiquitous across biological systems and are considered to contribute to the maintenance of cellular homeostasis. These adaptive responses activate protective mechanisms against diverse stressors, including physical challenges like radiation and temperature extremes, as well as chemical threats such as toxins, electrophiles, and reactive oxygen species to shield organisms from harm. Notably, Nrf2 activators like curcumin, resveratrol, and sulforaphane exert hormetic effects at low doses by acting as electrophiles that modify the regulatory protein Keap1 (15).

In this study, we also demonstrated that DNCB, a known inducer of the Nrf2 signaling pathway (9-11) exhibits an activating effect on metabolic activity at low doses. This effect was observed at the lower cell density (0.44 × 10^6^ cells/mL) across both tested exposure durations and reached statistical significance in a low-dose range between 0.05 µM and 0.5 µM DNCB (p<0.05). Additionally, a hormetic trend was observed at cell density of 1 × 10^6^ cells/mL. These findings indicate a reproducible hormetic response at low concentrations of DNCB. In this context, it should also be noted that, of the 29 individual experiments we analyzed in total, 15 showed a clear hormetic effect in the low-dose range (+10% to +63%), in 9 experiments no or only weak hormetic effects (+1% to +9%) was observed and in 5 experiments a negative effect (up to −26%) were observed (see Table S1). This further emphasizes the variability inherent in living systems, which must be considered in hormesis research.

Comparison with literature data indicates that the observed maximum stimulation (125%) and stimulation width (10-fold) are typical for hormetically active substances. In detail, Calabrese and Blain, 2011 showed that across 1,839 studies on metabolic endpoints, the most common stimulation maxima ranged from 110% to 150%, while in 1,213 studies on metabolic endpoints, the predominant stimulation widths ranged from 1-fold to 10-fold (2). Since DNCB triggers a common mechanism responsible for hormetic stimulation by activating the Nrf2 pathway (11, 16), it is consistent that DNCB exhibits stimulation magnitudes and ranges comparable to other hormetically active substances. Unlike most of the Nrf2 inducers studied, which are often secondary plant compounds, DNCB is a synthetically produced molecule. DNCB is of particular toxicological interest due to its extensive characterization over several decades. Interestingly, DNCB was also used in the 1990s to treat autoimmune diseases such as alopecia areata (17). Until a few years ago, the sensitizing and immune cell-activating potential of DNCB was also exploited in immunochemotherapy, for example in the treatment of skin metastases, and it is still mentioned as a treatment option (12) (13). Given the strong sensitizing, mutagenic (17), and genotoxic (18) properties of DNCB, which have long been recognized, its clinical relevance has declined substantially. With regard to its potential use in immunochemotherapy for skin metastases, the results of this study may be of interest, as we observed increased metabolic activity in the THP-1 tumor cell line at low doses. This raises the question of whether similar effects could also occur in other tumor cell types, which could be relevant with regard to a potential application in immunochemotherapy.

Due to its strong sensitizing potential, DNCB is primarily used in research, as a trigger substance for contact dermatitis in animal studies or as a positive control in toxicological assays evaluating the sensitizing potential of test compounds. It is also employed to induce atopic dermatitis in mouse models for testing novel therapeutic strategies (18). Studies have shown that a single dose of DNCB can suffice for sensitization, while repeated low doses over several days require a lower total amount to achieve the same effect (19). This implies that frequent, low-dose exposures pose a high sensitization risk in hazard assessments, with no established safe threshold currently identified for DNCB. The lowest concentrations tested in human studies, 60 µg (single application) or 10 µg (applied over three consecutive days), elicited sensitization in volunteers (19). The lowest DNCB concentrations reported in the literature for in vitro experiments are 1 µg/mL (equivalent to 5 µM) and 0.3 µg/mL (equivalent to 1.7 µM) (14). When using 5 µM DNCB on THP-1 cells (1 × 10^6^ cells/mL), Yoshida et al. observed a 50% increase in CD86 expression and a modest ~15% rise in CD54 expression, whereas at 1.7 µM, CD86 showed no change and CD54 exhibited a marginal 10% increase (14). These effects could not be replicated by Miyazawa et al., who found no increases in CD86 or CD54 at the lowest concentration tested (1.25 µg/mL eq. 6.38 µM) compared to the vehicle control (20). Our results with 1 × 10^6^ cells/mL also didn’t show an increased expression of CD86 and CD54 for 5µM DNCB exposure. Instead, using DNCB in lower concentration between 0.1 µM and 1 µM a maximum reduced expression level of −16% for CD86 and −12% for CD54 was observed. Hormetic responses are classically defined as compensatory or overcompensatory adaptations elicited by subtoxic perturbations of cellular homeostasis. In the present context, the reduction in CD86 and CD54 surface marker expression, and the consequent immunological hypo-responsiveness, does not constitute an overcompensatory response per se. Accordingly, it remains debatable whether this effect should be classified as a classically hormetic response. However, diminished surface expression of immune activation markers may also reflect a homeostatic adaptation. Immune activation is energetically demanding, and upon intracellular entry of a subtoxic compound, metabolic resources may be preferentially redirected toward rapid detoxification and elimination of the stressor. Only beyond a certain threshold may immune engagement become advantageous for maintaining organismal homeostasis. Thus, the observed mild hyporesponsiveness may represent a compensatory mechanism that preserves sufficient energy for essential cellular homeostatic processes. In this framework, the tendency toward hyporesponsiveness may likewise be interpreted as a hormetic effect.

Given that the most frequently reported stimulation maxima for the immunological endpoint lies between 110% and 150% relative to the control, the effect size observed here of −12% or −16% would be excepted. It is also noteworthy that the observed tendency toward hyporesponsiveness in the low-dose range suggests that a dose range exists in which DNCB does not elicit measurable sensitizing effects. However, it was not investigated whether this effect persists after repeated administration in the low-dose range, or whether the hyporesponsiveness eventually shifts to hyperresponsiveness. This would be entirely plausible, as the accumulation of perturbations may require an adaptive response to maintain homeostasis, a notion that is also supported by the results of in vivo studies (19).

To our knowledge, this study is the first to specifically investigate the low-dose range of DNCB and its potential effects. More comprehensive analyses beyond CD86 and CD54 would be required to draw definitive conclusions regarding immunological endpoints and their relevance for sensitizing potential, particularly in relation to in vivo conditions. The hormetic effect observed in metabolic activity may also be relevant for research on atopic dermatitis models and contact dermatitis mechanisms. Overall, these findings underscore the importance of incorporating low-dose ranges into chemical characterization and risk assessment.

### Limitations of the study

Although the hormetic effect reached statistical significance, not all individual experiments showed an increase in metabolic activity, underscoring the influence of variability in biological systems. This variability may be particularly pronounced in tumor cell lines, which often exhibit altered mitochondrial function, metabolic profiles, and heterogeneous mutational backgrounds.

## Acknowledgments

We thank Martin Thies for valuable assistance in measuring THP-1 cell respiration and Natalie Lobes for valuable assistance in measuring THP-1 cell respiration as well as in the analysis of cell surface markers.

## Author contributions

Darja Henseler: Conceptualization; project administration; data curation; formal analysis; visualization; writing original draft- and editing the manuscript.

Osmond A. Aruna: Data-Acquisition; formal analysis; statistics; visualization; writing-review and editing.

All authors consent to the final submitted manuscript.

## Material and methods

### Chemicals and Reagents

The chemicals 1-chlor-2,4-dinitro-benzene (DNCB) (CAS 97-00-7) and dimethyl sulfoxide (DMSO) (CAS 67-68-5) were purchased from Sigma Aldrich (Taufkirchen, Germany). Cell culture reagents RPMI 1640, DMEM, fetal bovine serum (FBS superior), 2-hydroxyethyl)piperazine-1-ethanesulfonic acid (HEPES), L-glutamine and antibiotic/antimycotic solution (10000 U/ml penicillin, 10 mg/ml streptomycin sulfate, 25 μg/ml amphotericin B) were obtained from Sigma Aldrich. The Invitrogen PrestoBlue™ HS Cell Viability Reagent was purchased from Thermo Fisher.

### Routine Cell Culture

THP-1 cells were obtained from the German Collection of Microorganisms and Cell Cultures GmbH (DSMZ, Braunschweig, Germany) and maintained in RPMI 1640 with phenol red supplemented with 10% (v/v) FBS superior, 25 mM HEPES, 4 mM L-glutamine, 1% antibiotic/antimycotic solution and 50 μM β-mercaptoethanol at 37°C, in a humidified atmosphere with 5% CO_2_. Cells were routinely split to 0.2 × 10^6^ cells/mL twice per week. Passage numbers 4 to 20 were used for the experiments.

### Determination of cell density

To determine which cell density is suitable for measuring metabolic activity, THP-1 cells were seeded in 96well-plates with cell densities ranging from 10,000 and 220,000 cells/well in a total volume of 180 µL of RPMI at 2 h and 24 h after seeding, 20 µL of PrestoBlue™ HS Cell Viability Reagent was added, followed by incubation for 30 min at 37°C in a humidified atmosphere containing 5% CO_2_. Fluorescence was measured using a fluorometer with excitation/emission filters of 530/25 and 590/35 nm respectively. The experiment was performed with 4 technical replicates.

### Chemical exposure of THP-1

For chemical exposure, THP-1 cells were seeded into 96 well-plates with a density of 80,000 or 180,000 cells/well in a volume of 90 µL exposure media (RPMI without phenol red, supplemented with HEPES, 10% FBS, and 1% L-glutamine). Chemicals were prepared as a serial dilution and added to the exposure medium at a ratio of 1:250. The chemical– exposure medium mixture was then added to the cells at a 1:1 ratio, resulting in a final volume of 180 µL per well. Final concentration of the chemicals was 0.05 µM, 0.1 µM, 0.5 µM, 1 µM, 5 µM, 20 µM and 40 µM DNCB and 0,2% DMSO as vehicle control. After chemical addition, the 96-well plate was sealed with foil and incubated for 4 h or 24 h, to investigate effects of short and longtime exposure. Each experiment was performed with six technical replicates. A total of 7 or 8 independent experiments were performed per condition with 80,000 cells (0.44 × 10^6^ cells/mL), and 8 or 9 independent experiments were performed per condition with 180,000 cells (1 × 10^6^ cells/mL). For analysis of cell surface markers CD86 and CD54, THP-1 cells were exposed to chemicals analogously, using an exposure time of 24 h. In this case, each experiment included four technical replicates, and 5 or 6 independent experiments were performed.

In the metabolic activity assay, a medium control was included in addition to the vehicle control to account for possible vehicle-related baseline effects. Relative to the medium control, the vehicle control exhibited moderate inter-experimental variability, with mean reductions of 20% at 80,000 cells and 10% at 180,000 cells. The coefficient of variation was between 18.6% and 21.2%. Data were normalized to the corresponding vehicle control (0.2% DMSO).

### Measurement of metabolic activity

The Invitrogen PrestoBlue™ HS Cell Viability Reagent was used to determine metabolic activity. The ready-to-use reagent a resazurin-based assay providing resazurin which is reduced to the fluorophore resorufin in metabolic active cells. Cells with higher metabolic activity exhibit increased conversion to resorufin and, consequently, higher fluorescence intensity. The reagent is non-toxic to cells and can be added directly to the culture for up to 24 h (according to the manufacturer’s instructions). To determine metabolic activity, 20 µL of the reagent was added afte 4 h and 24 h of chemical exposure (1:10 ratio), followed by gentle mixing and incubation for an additional 30 min at 37 °C and 5% CO_2_ without sealing the plate. Following incubation, fluorescence intensity was measured using a fluorometer with excitation and emission filters of 530/25 nm and 590/35 nm, respectively. THP-1 cells were directly analyzed in the 96-well plate. For data analysis, the blank signal (PrestoBlue™ and RPMI; 1:10) was subtracted from the measured fluorescence signal of PrestoBlue™-treated cells. As a quality control measure, the coefficient of variation (CV) was calculated for the technical replicates. In most cases, the CV ranged between 3% and 12%. In a few isolated experiments, the CV exceeded 20%. When this occurred in the solvent controls, the corresponding experiments were excluded from the analysis.

### Analysis of surfer markers CD86 and CD54

The expression of the cell surface markers CD86 and CD54 was quantified by flow cytometry using a BD FACSVerse™ system (BD Biosciences, Heidelberg, Germany). Following chemical exposure, THP-1 cells were collected by centrifugation (200 × g, 5 min, 20 °C) and washed twice with phosphate-buffered saline (PBS). Prior to antibody staining, cells were incubated in PBS containing bovine serum albumin (BSA) for 15 min at room temperature to block non-specific binding. Cells were then stained with fluorochrome-conjugated antibodies against CD86 (FITC, clone 2331 (FUN-1) and CD54 (FITC, clone 6.5B5), along with their respective isotype controls (Mouse IgG1/FITC). Antibodies were obtained from BD Biosciences (Heidelberg, Germany) and Dako (Jena, Germany).

The staining and analysis procedure was based on the human Cell Line Activation Test (h-CLAT) described in OECD Test Guideline 442E, with the following modifications: cells were exposed under the experimental conditions described above, including adjusted cell densities. In addition, a baseline expression threshold (MFI > 20 after isotype correction) was introduced as a quality control criterion. Fluorescence signals were acquired, and MFI values were determined. The relative expression of CD86 and CD54 was calculated by subtracting the MFI of the corresponding isotype control from the MFI of the specific antibody. A total of five independent experiments were performed, each including four technical replicates. For graphical visualization, MFI values of DNCB-treated samples were normalized to the corresponding vehicle control (0.2% DMSO). Only experiments fulfilling the predefined baseline expression criterion were included in the analysis to minimize the influence of background signal.

### Statistical Analysis

For the analysis of the metabolic endpoint, at least seven biological replicates per condition were included to increase statistical power. To achieve comparability of the signals across different experimental days, the mean fluorescence signal of each condition was normalized to the mean fluorescence signal of control (DMSO 0.2%). For statistical analysis, data were analyzed using GraphPad Prism version 10.6 (GraphPad Software, California, USA). All results are reported as mean percentages relative to the control (0.2% DMSO). An unpaired t-test with Welch’s correction was performed to compare DNCB-treated samples with the vehicle control (0.2% DMSO). A p value of <0.05 was considered to be statistically significant. For the analysis of immune endpoint at least five independent experiments were conducted, each with 4 technical replicates. Only experiments with a minimum MFI of 20 for vehicle control were accepted for analysis (baseline expression criterion). Data were visualised by using GraphPad Prism version 10.6 (GraphPad Software, California, USA). For statistical analysis, an unpaired t-test with Welch’s correction was performed to compare DNCB-treated samples with the control (0.2% DMSO). A p value of <0.05 was considered statistically significant.

## Supplementary Data

**Table S1.** Variability in metabolic effects across all experiments. A negative effect (a value below zero) is highlighted in yellow, whereas a weak hormetic effect (values between 1 and 9) is highlighted in light green and a hormetic effect of ≥10 is highlighted in dark green.

| Experimental condition | DNCB treatment [$\mu\text{M}$] | | | | | | |
| --- | --- | --- | --- | --- | --- | --- | --- |
|  | 0.05 | 0.1 | 0.5 | 1 | 5 | 20 | 40 |
|  | Metabolic activity (%) |  |  |  |  |  |  |
| 4 h, 80,000 cells | 96 | 89 | 93 | 91 | 92 | 78 | 84 |
| 4 h, 80,000 cells | 127 | 115 | 119 | 120 | 105 | 93 | 76 |
| 4 h, 80,000 cells | 130 | 131 | 116 | 117 | 115 | 127 | 67 |
| 4 h, 80,000 cells | 167 | 149 | 124 | 109 | 91 | 74 | 50 |
| 4 h, 80,000 cells | 159 | 163 | 140 | 113 | 95 | 67 | 49 |
| 4 h, 80,000 cells | 93 | 93 | 97 | 99 | 98 | 85 | 51 |
| 4 h, 80,000 cells | 110 | 87 | 110 | 90 | 87 | 95 | 61 |
| 4 h, 80,000 cells | 116 | 100 | 111 | 102 | 86 | 82 | 69 |
| 4 h, 180,000 cells | 142 | 132 | 123 | 130 | 111 | 99 | 72 |
| 4 h, 180,000 cells | 122 | 121 | 105 | 106 | 97 | 86 | 66 |
| 4 h, 180,000 cells | 111 | 100 | 105 | 107 | 97 | 89 | 92 |
| 4 h, 180,000 cells | 88 | 101 | 96 | 97 | 91 | 77 | 83 |
| 4 h, 180,000 cells | 105 | 105 | 92 | 92 | 96 | 86 | 97 |
| 4 h, 180,000 cells | 144 | 137 | 122 | 126 | 130 | 119 | 100 |
| 4 h, 180,000 cells | 91 | 93 | 91 | 109 | 93 | 85 | 79 |
| 24 h, 80,000 cells | 156 | 140 | 113 | 105 | 100 | 47 | 15 |
| 24 h, 80,000 cells | 101 | 99 | 101 | 100 | 104 | 45 | 18 |
| 24 h, 80,000 cells | 126 | 111 | 110 | 105 | 102 | 28 | 3 |
| 24 h, 80,000 cells | 140 | 129 | 112 | 107 | 77 | 29 | 8 |
| 24 h, 80,000 cells | 149 | 135 | 118 | 111 | 67 | 21 | 7 |
| 24 h, 80,000 cells | 101 | 97 | 94 | 92 | 93 | 23 | 4 |
| 24 h, 80,000 cells | 100 | 105 | 97 | 103 | 71 | 19 | 11 |
| 24 h, 180,000 cells | 126 | 114 | 112 | 111 | 113 | 54 | 8 |
| 24 h, 180,000 cells | 109 | 105 | 104 | 105 | 101 | 56 | 11 |
| 24 h, 180,000 cells | 115 | 114 | 107 | 110 | 108 | 49 | 15 |
| 24 h, 180,000 cells | 136 | 136 | 121 | 121 | 109 | 47 | 12 |
| 24 h, 180,000 cells | 100 | 103 | 100 | 100 | 101 | 51 | 19 |
| 24 h, 180,000 cells | 101 | 106 | 105 | 101 | 103 | 56 | 5 |
| 24 h, 180,000 cells | 96 | 95 | 92 | 93 | 109 | 57 | 14 |

**Table S2.** Vehicle control in comparison to medium. Shown are the metabolic activities, expressed as percentages of the vehicle control (0.2% DMSO) relative to the RPMI control, together with the calculated coefficients of variation (CVs).

| Experimental condition | DMSO 0.2 % |  |  |  |
| --- | --- | --- | --- | --- |
|  | 4 h<br>80,000 cells | 4 h<br>180,000 cells | 24 h<br>80,000 cells | 24 h<br>180,000 cells |
| Metabolic activity compared to medium control (%) | 97 | 71 | 55 | 76 |
|  | 77 | 72 | 93 | 87 |
|  | 71 | 74 | 82 | 80 |
|  | 52 | 96 | 69 | 62 |
|  | 56 | 105 | 66 | 115 |
|  | 93 | 64 | 92 | 100 |
|  | 96 | 108 | 99 | 101 |
|  | 89 | - | - | - |
| CV (%) | 21,2 | 19,9 | 19,1 | 18,6 |

**Table S3.** Overview of all planned and performed experiments assessing metabolic activity after DNCB exposure using 80,000 cells.

| Treatment | Fluorescence Intensity after 4 h exposure (80,000 cells) |  |  |  |  |  |  |  |  |  |
| --- | --- | --- | --- | --- | --- | --- | --- | --- | --- | --- |
|  | Exp1 | Exp2 | Exp3 | Exp4 | Exp5 | Exp6 | Exp7 | Exp8 | Mean |  |
| Basal Medium | 2060 | 2623 | 1865 | 2313 | 2658 | 3055 | 2692 | 1683 | 2369 |  |
| Control (DMSO 0.2%) | 1989 | 2025 | 1320 | 1202 | 1480 | 2855 | 2574 | 1496 | 1867 |  |
| DNCB 0.05 μM | 1904 | 2569 | 1717 | 2009 | 2346 | 2641 | 2833 | 1736 | 2219 |  |
| DNCB 0.1 μM | 1765 | 2332 | 1728 | 1788 | 2409 | 2651 | 2234 | 1498 | 2051 |  |
| DNCB 0.5 μM | 1849 | 2419 | 1534 | 1485 | 2079 | 2760 | 2833 | 1654 | 2076 |  |
| DNCB 1 μM | 1814 | 2432 | 1542 | 1313 | 1678 | 2817 | 2326 | 1527 | 1931 |  |
| DNCB 5 μM | 1839 | 2121 | 1524 | 1090 | 1400 | 2795 | 2242 | 1282 | 1787 |  |
| DNCB 20 μM | 1551 | 1881 | 1680 | 892 | 994 | 2434 | 2443 | 1223 | 1637 |  |
| DNCB 40 μM | 1664 | 1544 | 886 | 605 | 728 | 1457 | 1570 | 1025 | 1185 |  |
| Treatment | Ratio to control (%) after 4 h exposure (80,000 cells) |  |  |  |  |  |  |  |  | p-value |
|  | Exp1 | Exp2 | Exp3 | Exp4 | Exp5 | Exp6 | Exp7 | Exp8 | Mean |  |
| DNCB 0.05 μM | 96 | 127 | 130 | 167 | 158 | 93 | 110 | 116 | 125 | 0.0321 |
| DNCB 0.1 μM | 89 | 115 | 131 | 149 | 163 | 93 | 87 | 100 | 116 | 0.1488 |
| DNCB 0.5 μM | 93 | 119 | 116 | 124 | 140 | 97 | 110 | 111 | 114 | 0.0286 |
| DNCB 1 μM | 91 | 120 | 117 | 109 | 113 | 99 | 90 | 102 | 105 | 0.1898 |
| DNCB 5 μM | 92 | 105 | 116 | 91 | 95 | 98 | 87 | 86 | 96 | 0.3968 |
| DNCB 20 μM | 78 | 93 | 127* | 74 | 67 | 85 | 95 | 82 | 82 | 0.1176 |
| DNCB 40 μM | 84 | 76 | 67 | 50 | 49 | 51 | 61 | 69 | 63 | 0.0004 |
| Included | yes | yes | yes | yes | yes | yes | yes | yes |  |  |
| Treatment | Fluorescence Intensity after 24 h exposure (80,000 cells) |  |  |  |  |  |  |  |  |  |
|  | Exp1 | Exp2 | Exp3 | Exp4 | Exp5 | Exp6 | Exp7 | Mean |  |  |
| Basal Medium | 4954 | 6108 | 3606 | 3896 | 5057 | 4588 | 4895 | 4729 |  |  |
| Control (DMSO 0.2%) | 2702 | 5672 | 2967 | 2681 | 3347 | 4234 | 4865 | 3781 |  |  |
| DNCB 0.05 μM | 4226 | 5707 | 3753 | 3751 | 4999 | 4261 | 4867 | 4509 |  |  |
| DNCB 0.1 μM | 3782 | 5620 | 3308 | 3469 | 4505 | 4097 | 5095 | 4268 |  |  |
| DNCB 0.5 μM | 3048 | 5702 | 3254 | 3013 | 3957 | 3984 | 4725 | 3955 |  |  |
| DNCB 1 μM | 2846 | 5672 | 3124 | 2859 | 3720 | 3898 | 5002 | 3874 |  |  |
| DNCB 5 μM | 2707 | 5913 | 3035 | 2075 | 2255 | 3918 | 3449 | 3336 |  |  |
| DNCB 20 μM | 1276 | 2562 | 830 | 775 | 718 | 983 | 946 | 1156 |  |  |
| DNCB 40 μM | 400 | 1025 | 79 | 221 | 239 | 163 | 555 | 383 |  |  |
| Treatment | Ratio to control (%) after 24 h exposure (80,000 cells) |  |  |  |  |  |  |  | p-value |  |
|  | Exp1 | Exp2 | Exp3 | Exp4 | Exp5 | Exp6 | Exp7 | Mean |  |  |
| DNCB 0.05 μM | 156 | 101 | 127 | 140 | 149 | 101 | 100 | 125 | 0.0324 |  |
| DNCB 0.1 μM | 140 | 99 | 111 | 129 | 135 | 97 | 105 | 117 | 0.0423 |  |
| DNCB 0.5 μM | 113 | 101 | 110 | 112 | 118 | 94 | 93 | 106 | 0.0825 |  |
| DNCB 1 μM | 105 | 100 | 105 | 107 | 111 | 92 | 106 | 104 | 0.1302 |  |
| DNCB 5 μM | 100 | 104 | 102 | 77 | 67 | 93 | 69 | 88 | 0.0985 |  |
| DNCB 20 μM | 47 | 45 | 28 | 29 | 21 | 23 | 27 | 32 | <0.0001 |  |
| DNCB 40 μM | 15 | 18 | 3 | 8 | 7 | 4 | 59 | 16 | <0.0001 |  |
| Included | yes | yes | yes | yes | yes | yes | yes |  |  |  |

Red numbers indicate a coefficient of variation >20.
\* Value was excluded from analysis because it is considered as an outlier (lies more than 2 or 3 standard deviations from the mean).

**Table S4.**
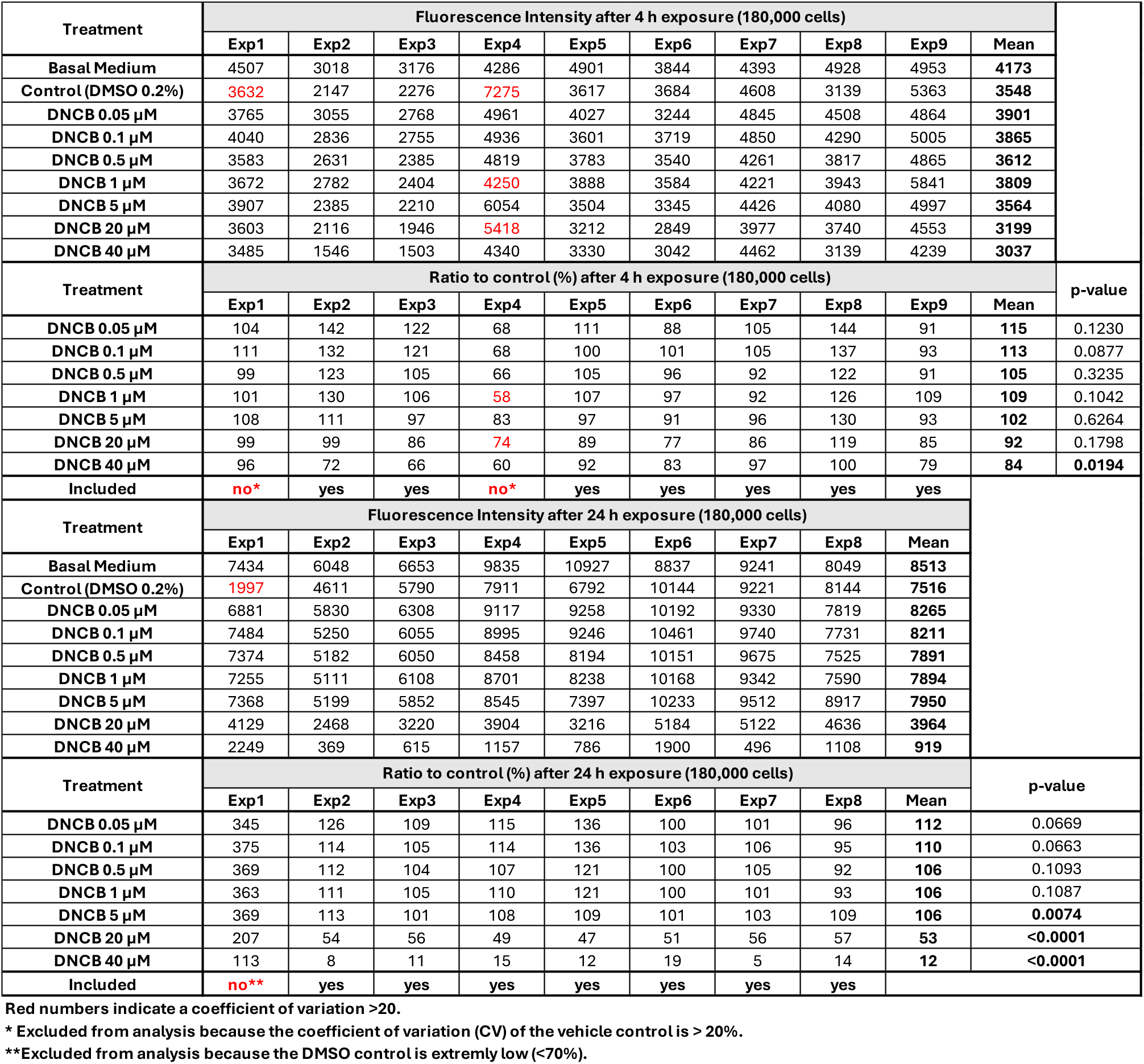
Overview of all planned and performed experiments assessing metabolic activity after DNCB exposure using 180,000 cells.

**Table S5.** Overview of all planned and performed experiments analyzing CD86 and CD54 expression using 80,000 cells.

| Treatment | CD86 expression (ratio to control %) on THP-1 cells (80,000 cells) |  |  |  |  |  |
| --- | --- | --- | --- | --- | --- | --- |
| DNCB [μM] | Exp 1 | Exp 2 | Exp 3 | Exp 4 | Exp 5 | Mean |
| 0.05 | n.a. | 85% | 93% | 102% | 109% | 97% |
| 0.1 | n.a. | 82% | 79% | 95% | 104% | 90% |
| 0.5 | n.a. | 73% | 72% | 94% | 95% | 84% |
| 1 | n.a. | 69% | 84% | 89% | 94% | 84% |
| 5 | n.a. | 114% | 97% | 165% | 137% | 128% |
| 20 | n.a. | 120% | 209% | 123% | 144% | 149% |
| 40 | n.a. | 153% | 80% | 67% | 97% | 99% |
| MFI of vehicle control | 13 | 24 | 21 | 29 | 21 |  |
| Included | no* | yes | yes | yes | no* |  |
| Treatment | CD54 expression (ratio to control %) on THP-1 cells (80,000 cells) |  |  |  |  |  |
| DNCB [μM] | Exp 1 | Exp 2 | Exp 3 | Exp 4 | Exp 5 | Mean |
| 0.05 | n.a. | 105% | n.a. | 93% | 89% | 96% |
| 0.1 | n.a. | 94% | n.a. | 91% | 95% | 93% |
| 0.5 | n.a. | 102% | n.a. | 94% | 90% | 95% |
| 1 | n.a. | 94% | n.a. | 82% | 87% | 88% |
| 5 | n.a. | 152% | n.a. | 224% | 135% | 170% |
| 20 | n.a. | 185% | n.a. | 132% | 98% | 138% |
| 40 | n.a. | 52% | n.a. | 22% | 36% | 36% |
| MFI of vehicle control | 15 | 35 | 17 | 63 | 81 |  |
| Included | no* | yes | no* | yes | yes |  |

\*Excluded from analysis because the MFI of the vehicle control was <20.

**Table S6.** Overview of all planned and performed experiments analyzing CD86 and CD54 expression using 180,000 cells.

| Treatment | CD86 expression (ratio to control %) on THP-1 cells (180,000 cells) |  |  |  |  |  |  |
| --- | --- | --- | --- | --- | --- | --- | --- |
| DNCB [μM] | Exp 1 | Exp 2 | Exp 3 | Exp 4 | Exp 5 | Exp 6 | Mean |
| 0.05 | 102% | 86% | 90% | n.a. | 104% | n.a. | 95% |
| 0.1 | 103% | 101% | 83% | n.a. | 79% | n.a. | 92% |
| 0.5 | 109% | 86% | 73% | n.a. | 80% | n.a. | 87% |
| 1 | 101% | 79% | 75% | n.a. | 81% | n.a. | 84% |
| 5 | 106% | 108% | 82% | n.a. | 100% | n.a. | 99% |
| 20 | 280% | 311% | 218% | n.a. | 364% | n.a. | 293% |
| 40 | 65% | 69% | 57% | n.a. | 71% | n.a. | 66% |
| MFI of vehicle control | 30 | 26 | 28 | 12 | 21 | 18 |  |
| Included | yes | yes | yes | no* | yes | no* |  |
| Treatment | CD54 expression (ratio to control %) on THP-1 cells (180,000 cells) |  |  |  |  |  |  |
| DNCB [μM] | Exp 1 | Exp 2 | Exp 3 | Exp 4 | Exp 5 | Exp 6 | Mean |
| 0.05 | 103% | 100% | 103% | n.a. | 94% | 99% | 100% |
| 0.1 | 95% | 110% | 113% | n.a. | 62% | 118% | 99% |
| 0.5 | 98% | 101% | 95% | n.a. | 61% | 103% | 91% |
| 1 | 94% | 96% | 94% | n.a. | 69% | 96% | 90% |
| 5 | 80% | 92% | 85% | n.a. | 85% | 96% | 88% |
| 20 | 388% | 591% | 523% | n.a. | 458% | 422% | 477% |
| 40 | 26% | 54% | 32% | n.a. | 68% | 43% | 45% |
| MFI of vehicle control | 61 | 37 | 42 | 15 | 21 | 36 |  |
| Included | yes | yes | yes | no* | yes | yes |  |

\*Excluded from analysis because the MFI of the vehicle control was <20.

